# Shoot-derived miR2111 controls legume root and nodule development

**DOI:** 10.1101/2020.09.02.278739

**Authors:** Mengbai Zhang, Huanan Su, Peter M. Gresshoff, Brett J. Ferguson

## Abstract

Legumes control their nodule numbers through the Autoregulation Of Nodulation (AON). Rhizobia infection stimulates the production of root-derived CLE peptide hormones that are translocated to the shoot where they regulate a new signal. We used soybean to demonstrate that this shoot-derived signal is miR2111, which is transported via phloem to the root where it targets transcripts of *Too Much Love* (*TML*), a negative regulator of nodulation. Shoot perception of rhizobia-induced CLE peptides suppresses miR2111 expression, resulting in TML accumulation in roots and subsequent inhibition of nodule organogenesis. Feeding synthetic mature miR2111 via the petiole increased nodule numbers per plant. Likewise, elevating miR2111 availability by over-expression promoted nodulation, while target mimicry of *TML* induced the opposite effect on nodule development in wild-type plants and alleviated the supernodulating and stunted root growth phenotypes of AON-defective mutants. Additionally, in non-nodulating wild-type plants, ectopic expression of miR2111 significantly enhanced lateral root emergence with a decrease in lateral root length and average root diameter. In contrast, hairy roots constitutively expressing the target mimic construct exhibited reduced lateral root density. Overall, these findings demonstrate that miR2111 is both the critical shoot-to-root factor that positively regulates root nodule development, and also acts to shape root system architecture via orchestrating the degree of root branching, as well as the length and thickness of lateral roots.

## Introduction

Rhizobia bacteria can convert atmospheric di-nitrogen into compounds plants can use. They do so inside specialised nodule structures that form on the roots of legume host plants, making this symbiosis one of the world’s most beneficial plant-microbe interactions (Ferguson *et al*., 2010; Oldroyd, 2013). Forming and maintaining nodules is resource intensive and hence the host plant tightly regulates the number of nodules it forms (Soyano and Kawaguchi, 2014; Ferguson *et al*., 2019). Understanding the molecular components that regulate legume nodule numbers is viewed as pivotal to efforts aimed at optimising nitrogen fixation in agriculture.

One of the key mechanisms driving legume nodulation control is called Autoregulation Of Nodulation (AON; Caetano-Anollés and Gresshoff, 1991). It is a systemic process, initiated by the production of tri-arabinosylated CLE peptides in the host root following infection with compatible rhizobia bacteria (Okamoto *et al*., 2013) and therefore only activated in rhizobia-inoculated legume plants. These peptides are transported in xylem to the shoot where they are perceived by a Leucine-Rich-Repeat Receptor-Kinase (LRR-RK; GmNARK/PvNARK/LjHAR1/MtSUNN1/PsSYM29; Sagan and Duc, 1996; Krusell *et al*., 2002; Searle *et al*., 2003; Schnabel *et al*., 2005; Ferguson *et al*., 2014), likely in complex with other transmembrane factors (Miyazawa *et al*., 2010; Krusell *et al*., 2011; Crook *et al*., 2016). This receptor complex is located on the extracellular surface of leaf phloem parenchyma cells (Nontachaiyapoom *et al*., 2007). Perception of the CLE peptides leads to the regulation of a new factor in the shoot (recently proposed to be miR2111 in *Lotus japonicus*; Tsikou *et al*., 2018), which is subsequently transported to the roots where it regulates continued nodule organogenesis. Genetic defects in the AON molecular network can result in excessive nodule formation, which is referred to as hyper- or super-nodulation (Sagan and Duc, 1996; Krusell *et al*., 2002; Searle *et al*., 2003; Schnabel *et al*., 2005; Magori *et al*., 2009; Miyazawa *et al*., 2010; Krusell *et al*., 2011; Ferguson *et al*., 2014; Crook *et al*., 2016). Interestingly, these LRR-RK mutants also tend to exhibit altered root development in the absence of rhizobia, both in regard to lateral root numbers or root length (Day *et al*., 1986; Sagan and Duc, 1996; Wopereis *et al*., 2000; Schnabel *et al*., 2005).

An additional root factor known to regulate legume nodule numbers is a nucleus-localised Kelch repeat-containing F-box protein identified in *L. japonicus*, called Too Much Love (TML) (Takahara *et al*., 2013). As a subunit of an E3 ubiquitin complex, TML binds specifically to its protein substrate(s) via its Kelch-repeat sequence to mark the substrate for proteolytic degradation (Takahara *et al*., 2013). Mutations in *LjTML* cause hypernodulation and it was shown to function as a direct component in the AON pathway (Takahara *et al*., 2013; Soyano *et al*., 2014). Interestingly, mRNA transcripts of the closest orthologues of LjTML in *Arabidopsis thaliana* and soybean (*Glycine max*) are known to be targeted and cleaved by miR2111 (Hsieh *et al*., 2009; Pant *et al*., 2009; Xu *et al*., 2013; Zhang *et al*., 2014). This microRNA was initially reported to be induced by phosphate starvation, with its expression reduced within three-hours of Pi-replenishment (Hsieh *et al*., 2009; Pant *et al*., 2009). Previous studies have also identified miR2111 in phloem sap of *Brassica napus* and *Cucumis sativus* (Pant *et al*., 2009; Zhang *et al*., 2016). Therefore, it seemed highly plausible that miR2111 could be the shoot-derived signal of AON, produced in the shoot and transported in the phloem to the root where it regulates TML transcript levels to control legume root nodulation.

The critical shoot-derived factor of AON remained highly elusive since it was first predicted in the 1980s from a range of mutant- and physiological-based studies (Kosslak and Bohlool, 1984; Carroll *et al*., 1985; Delves *et al*., 1986). We used the commercially significant legume crop, soybean, to demonstrate this factor is miR2111. These findings are consistent with those recently reported using the model legume *L. japonicus* (Tsikou *et al*., 2018). We further demonstrate that the shoot**-**derived miR2111 signal has a role in regulating root system architecture.

Following production in the shoot, miR2111 is transported in the phloem to the root where it targets mRNA transcripts of TML for degradation. In AON, perception of the rhizobia-induced CLE peptides in the shoot results in a downregulation of miR2111, and hence an increased abundance of TML in the root, preventing continued nodule formation. Manipulation of miR2111 abundance by over-expression, feeding of exogenous miRNA, or target mimicry all resulted in a considerable change in nodule numbers. Moreover, in uninoculated plants, constitutive expression of miR2111 significantly enhanced root branching with reduced lateral root elongation and thickening, whereas transgenic expression of *TML* target-mimics decreased lateral root density. Findings from our studies to discover and functionally characterise miR2111 as the shoot-derived mobile signal of root and nodule development are reported here.

## Results

### miR2111 levels following rhizobia inoculation and the complete miR2111-encoding gene family of soybean

To establish whether miR2111 is regulated following rhizobia inoculation, stem-loop RT-PCR was used to quantify mature miR2111 levels in shoots and roots of *G. max* plants. Primers were specifically designed to detect both the canonical isoform and other variants of miR2111. Inoculation triggered a significant downregulation of miR2111 in leaves and roots of wild-type plants (∼85%), whereas only a moderate decrease (∼45%) was observed in leaves of a supernodulating *Gmnark* non-sense mutant (called *nts382*), indicating miR2111 expression is indeed controlled downstream of GmNARK in the AON pathway (Figure 1A).

**Figure 1.**
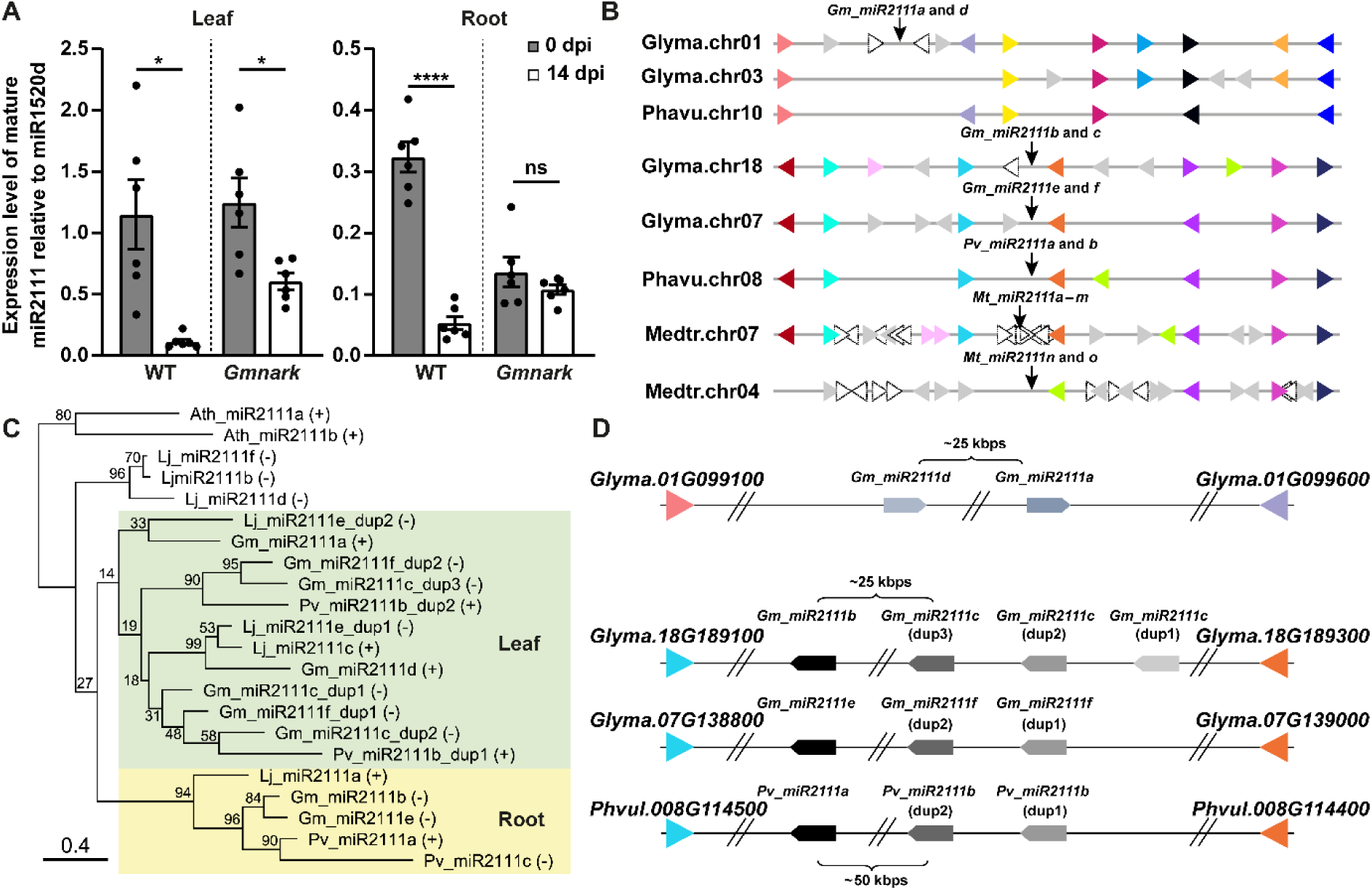
miR2111 family members from different species and expression of mature miR2111 in soybean following rhizobia inoculation. **(A)** Mature miR2111 levels in wild-type (WT) and *Gmnark* supernodulating soybean leaves and roots at 0 and 14 days post inoculation (dpi) with *Bradyrhizobium diazoefficiens*. Data are shown as mean ± SEM (*n* = 6 where each biological replicate was pooled from three individual plants grown in different pots). Asterisks denote statistically significant differences; \**p* < 0.05, \*\*\*\**p* < 0.0001 (Student’s *t*-test); ns, not significant. **(B)** Genomic microsynteny of *miR2111* family members of soybean, common bean and *M. truncatula*. Each triangle symbolises a gene and its orientation. Dotted white triangles are hypothetical genes with unknown functions. Genes belonging to the same family are coloured the same, with all other genes shown in grey. The genomic loci of the *miR2111* members are aligned with respect to the neighbouring genes within the *G. max, P. vulgaris* and *M. truncatula* genomes. **(C)** A phylogenetic tree of twenty-two miR2111 precursor sequences identified in *A. thaliana, G. max, L. japonicus* and *P. vulgaris*. **(D)** Genomic arrangement of soybean and common bean *miR2111* members in the context of their microsyntenies presented in **(B)**. Identically shaded arrowed bands represent closest orthologous pre-miR2111 sequences. See also Supplemental Figure 1 and Supplemental Table 1.

To determine which *miR2111* genes are regulated during AON, the entire gene family first had to be identified and characterised. In soybean, there are six genes encoding for mature miR2111 products. The canonical variant confirmed by small RNA sequencing studies in other species (Hsieh *et al*., 2009; Pant *et al*., 2009; Zhang *et al*., 2016), is generated from seven microRNA precursors of five *Gm-miR2111* genes, whereas two rare isoforms can be produced from one out of nine pre-miR2111 sequences (Supplemental Figure 1A). Six *miR2111* genes are distributed across three chromosomes, with *miR2111a* and *d, b* and *c*, and *e* and *f* located on Chromosomes 1, 18 and 7, respectively (Figure 1B and 1D). Each is located ∼25 kb from the other miR2111-encoding gene on that chromosome. Amongst them, *miR2111c* and *miR2111f* consist of multiple tandemly duplicated microRNA precursors, with individual duplicates referred to here as dup1, dup2, *etc*., based on their order within the locus (Figure 1C and 1D). Similarly, in *Phaseolus vulgaris* (common bean), *Pv-miR2111a* and *b* are separated by ∼50 kb on the same chromosome (Chr8) and the b locus comprises two pre-miR2111 duplicates in tandem (Figure 1B and 1D). Interestingly, duplication of pre-miR2111 is conserved in legumes, occurring in one *L. japonicus* and six *Medicago truncatula* miR2111 loci (Figure 1C and 1D; Supplemental Figure 1A and 1B; Supplemental Table 1). The precursor duplicates of *Gm-miR2111c* and *f*, and *Pv-miR2111b*, are only interspaced by ∼90 bp and were demonstrated to be expressed as single transcript with a shared promoter by mapping of overlapping RNAseq reads and subsequent cDNA amplification (Figure 1D and Supplemental Figure 2). Interestingly, *Pv-miR2111c* does not appear to have an orthologous copy in soybean, and in fact is located at the same locus as another protein-encoding gene (*Phvul*.*010G072200*), but on the opposite DNA strand with its own promoter (Supplemental Figure 1A).

High conservation is primarily seen between pre-miR2111 sequences across multiple species, barring loop regions (Supplemental Figure 1A and 1B). A highly conserved motif (TATAAT/TATAAAT) located ∼100-150 bp upstream of the pre-miR2111 sequence likely corresponds to the TATA box (Supplemental Figure 1A and 1B). This is supported by RNAseq findings where no reads can be mapped to the adjacent region beyond the identified motif (Supplemental Figure 2) and the 5’ RACE analysis of *Lj-miR2111* genes (Tsikou *et al*., 2018).

A phylogenetic tree based on the multiple sequence alignment presented in Supplemental Figure 1A revealed that *Gm-miR2111b* and *e* are likely duplicates (i.e. homeologues) with *Pv-miR2111a* as their likely orthologue (Figure 1C and 1D). Likewise, *Gm-miR2111c* and *f* appear to be duplicates with *Pv-miR2111b* as their orthologue (Figure 1C and 1D). In contrast, a lack of conserved pre-miR2111 sequences in the *G. max* Chromosome 3 region displaying similar syntenies of *Gm-miR2111a* and *d* may indicate that they are descended from the gene progenitors of *Gm-miR2111c* and *f* after the most recent genome-wide duplication of soybean ∼13 MYA (Figure 1B) (Schmutz *et al*., 2010). Alternatively, chromosomal rearrangement or deletions may have resulted in a loss of two miR2111 genes on *G. max* Chromosome 3. Being unique to common bean, *Pv-miR2111c* seems to have evolved from *Pv-miR2111a* after the divergence from soybean ∼19 MYA (McClean *et al*., 2010).

### Expression of miR2111-encoding genes and their regulation following rhizobia inoculation

To determine which miR2111-encoding genes are contributing to the pool of mature miR2111 in roots and leaves, including those being regulated by AON, the expression of each gene was assessed in soybean following rhizobia inoculation. Expression of *miR2111a, c, d* and *f* was mainly in the leaf, whereas *miR2111b* and *e* was predominately in the root (Figure 2). Therefore, the extremely low transcript abundance of *miR2111b* and *e* in the leaf and *miR21111a, c, d* and *f* in the root indicates that the contribution of these transcripts to the overall level of mature miR2111 is likely negligible. The transcript abundance of all miR2111-encoding genes decreased in wild-type soybean leaves following *B. diazoefficiens* inoculation, with most exhibiting a significant reduction in expression (Figure 2A). In contrast, almost no reduction in the miR2111-encoding genes was observed in roots of the same plants (Figure 2C). This indicates that the decline of mature miR2111 detected in the leaf and root (Figure 1A) is caused by a reduction in the expression of miR2111-encoding genes in the leaf (Figure 2A). Similar results were achieved using RNAseq (Supplemental Figure 2), where inoculation with compatible *B. diazoefficiens* resulted in the decreased expression of miR2111-encoding genes in the leaf, which was not observed in plants inoculated with an incompatible Nod Factor synthesis mutant (*NodC*^*-*^).

**Figure 2.**
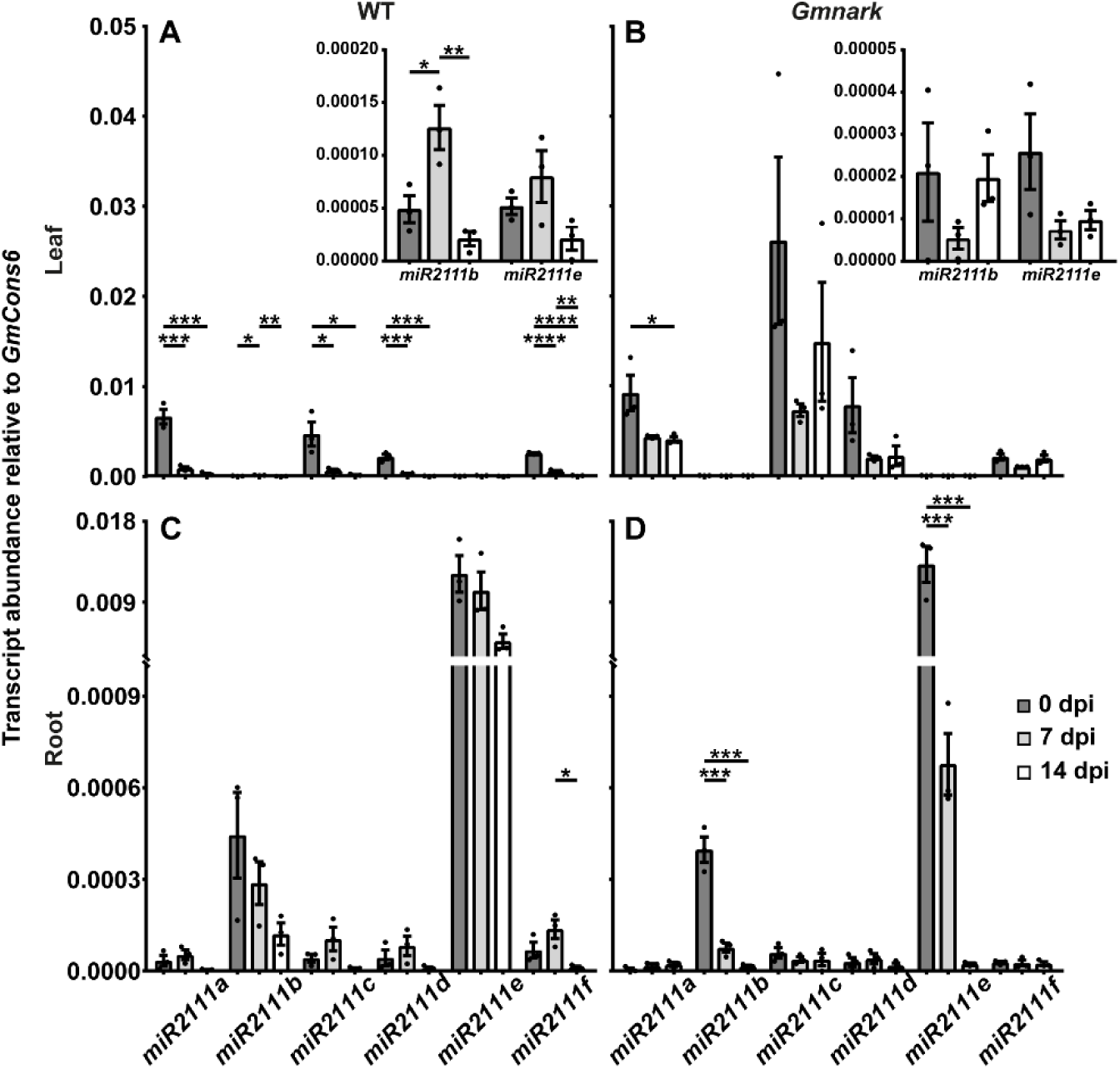
Transcript abundance of pri-miR2111 members in inoculated and uninoculated soybean plants. **(A-D)** Abundance of *Gm-miR2111a, b, c, d, e* and *f* primary transcripts in mature trifoliate leaves and roots of wild-type **(A and C)** or *Gmnark* **(B and D)** supernodulating plants. Data are shown as mean ± SEM (*n* = 3 where each biological replicate was pooled from three individual plants grown in different pots). Asterisks denote statistically significant differences; \**p* < 0.05, \*\**p* < 0.01, \*\*\**p* < 0.001, \*\*\*\**p* < 0.0001 (One-way ANOVA with post-hoc Tukey HSD test). See also Supplemental Figure 2.

In *Gmnark* mutant plants, which cannot detect the rhizobia-induced CLE peptides and hence are not predicted to regulate the shoot-derived factor of AON, miR2111 transcript levels were not suppressed in leaves to the same extent as observed in wild-type plants (Figure 2A and 2B). In particular, the significant downregulation of *miR2111c, d*, and *f* in wild-type leaves is not observed in the mutant (Figure 2A and 2B). This is consistent with our finding that leaves of *Gmnark* mutant plants did not exhibit as strong of a reduction of mature miR2111 compared with wild-type plants following *B. diazoefficiens* inoculation (Figure 1A). Interestingly, *miR2111b* and *e* were significantly reduced in nodulating *Gmnark* mutant roots, which may be a compensatory response to the significantly high level of miR2111 coming from the shoot compared with inoculated wild-type plants (Figure 2B and 2D). Despite some changes in the expression of miR2111-encoding genes in *Gmnark* mutant plants, there was no difference in the level of mature miR2111 in the roots (Figure 1A). These data demonstrate that the regulation of leaf-specialised miR2111-encoding genes occurs downstream of GmNARK in soybean.

### Level of miR2111 in sap and petiole-feeding synthetic double-stranded miR2111

To establish whether the level of mature miR2111 is also decreased in phloem sap following nodule organogenesis, sap was collected from stem tissue of non-inoculated and 14 day-post-inoculated wild-type soybean plants. Using absolute stem-loop RT-PCR, mature miR2111 was found to be significantly reduced in sap following rhizobia inoculation (Figure 3A). This is consistent with the reduced expression of miR2111-encoding genes in leaves (Figure 2A), and the reduced level of mature miR2111 in roots (Figure 1A) of inoculated wild-type plants.

**Figure 3.**
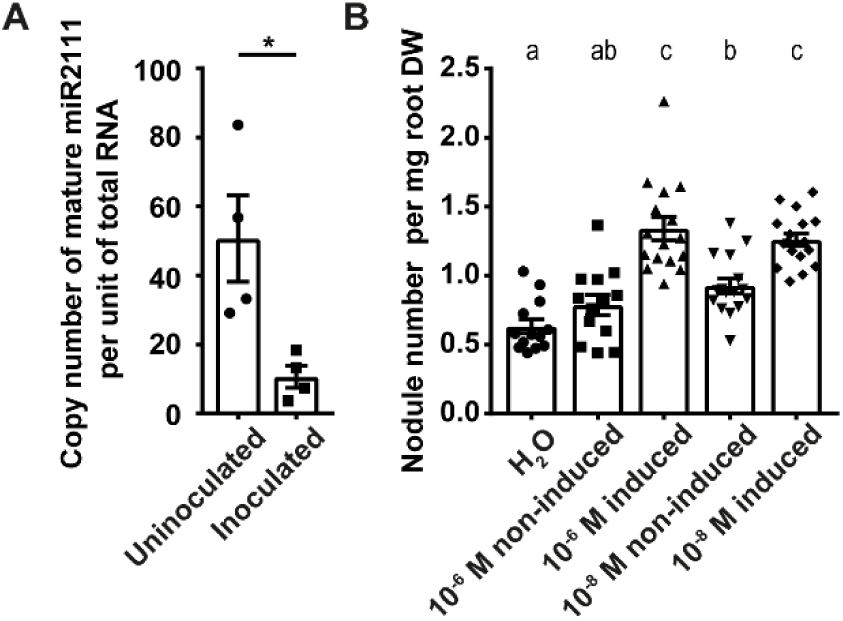
Abundance of mature miR2111 in soybean phloem sap and nodule number of soybean plants fed with mature miR2111. **(A)** Mature miR2111 level in phloem sap of wild-type soybean plants inoculated with or without *B. diazoefficiens*. Data are shown as mean ± SEM (*n* = 4, with 20-25 pooled plants per biological replicate). Asterisks indicate significant differences; \**p* < 0.05 (Student’s *t*-test). **(B)** Normalised nodule number of WT soybean plants treated via petiole feeding with induced (+) and non-induced (-) double-stranded RNA extracts containing 10^−6^ or 10^−8^ M miR2111 (induced), an equally diluted extract background (non-induced), or water (control). Data are shown as mean ± SEM (*n* ≥ 13). Different letters indicate statistical significance (One-way ANOVA with post-hoc Tukey HSD test). The experiments were repeated twice **(A)** and three times **(B)** independently with similar results. See also Supplemental Figures 3 and 4.

Characterising the role of miR2111 in nodule formation was achieved by synthesising mature miR2111 and feeding it to recipient soybean plants. Double-stranded miR2111 was synthesised using a bacterial IPTG-inducible expression system (Baneyx, 1999). The mature miR2111 product was subsequently collected and continuously fed into the vasculature of wild-type soybean plants via the petiole (Lin *et al*., 2011) to simulate miR2111 production from the leaf. Feeding both 10^−6^ and 10^−8^ M miR2111 treatments markedly raised the nodule number of recipient plants compared with non-induced samples or water control (Figure 3B; Supplemental Figures 3 and 4). This demonstrates a positive role for miR2111 in regulating legume nodulation.

**Figure 4.**
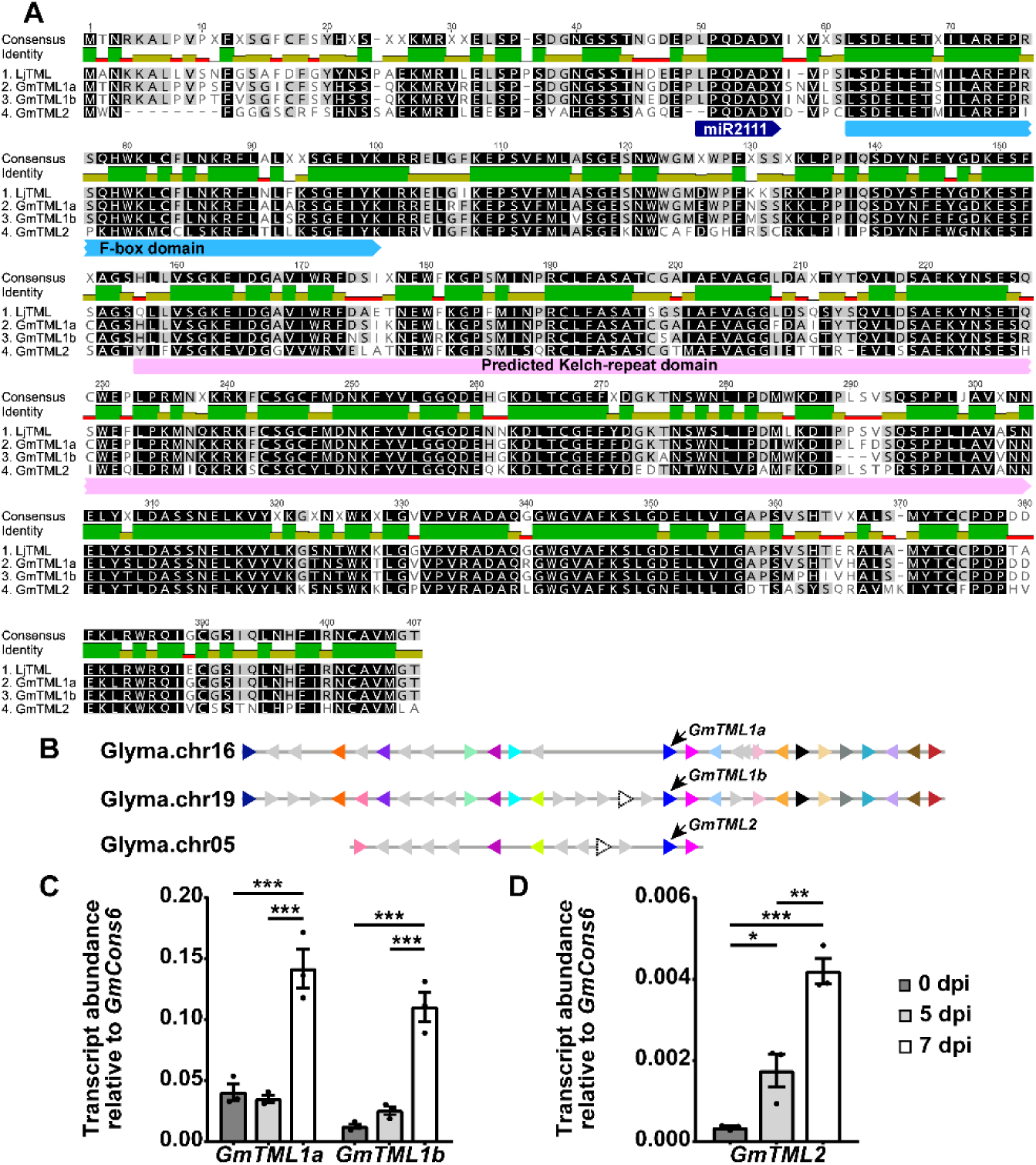
TML family members of soybean and their transcript abundance in roots following inoculation with *B. diazoefficiens*. **(A)** Protein sequence alignment of TML members, with miR2111 target sites (dark blue arrows), F-box domains (light blue arrows) and Kelch-repeat domains (pink arrows) shown. The darker the shading of each amino acid, the more highly conserved it is amongst the different family members. **(B)** Genomic environment of TML members of soybean showing microsynteny. **(C and D)** Transcript abundance of *GmTML1a* and *b* **(C)** or *GmTML2* **(D)** in spot-inoculated soybean roots. Data are shown as mean ± SEM (*n* = 3 where each biological replicate was pooled from three individual plants grown in different growth pouches). Asterisks indicate significant differences; \**p* < 0.05, \*\**p* < 0.01, \*\*\**p* < 0.001 (One-way ANOVA with post-hoc Tukey HSD test). See also Supplemental Figure 5.

### Transcriptional and bioinformatic analysis of the miR2111 targets, the TML family members

Previous work using soybean and *Arabidopsis* demonstrated that miR2111 targets mRNA transcripts of Kelch repeat-containing F-Box proteins (Hsieh *et al*., 2009; Pant *et al*., 2009; Xu *et al*., 2013; Zhang *et al*., 2014). Mutations in one of these proteins from *L. japonicus*, called Too Much Love, results in a hyper-nodulation phenotype (Magori *et al*., 2009; Takahara *et al*., 2013). Three soybean genes encoding products showing strong sequence homology with LjTML (AK339024) and exhibiting putative miR2111 cleavage sites were identified here (Figure 4A). With approximately 85% amino acid identity, GmTML1a (Glyma.16G057300) and GmTML1b (Glyma.19G090600), which are homeologous duplicates, are the closest orthologues to LjTML, whereas the pairwise identity between GmTML2 (Glyma.05G077700) and LjTML is just above 65%. *GmTML2* has a comparatively longer stretch of microcolinearity with *GmTML1b*, and likely originates from an ancient segmental duplication of the chromosome region containing *GmTML1b* (Figure 4B).

Spot inoculation of soybean seedlings grown in growth pouches (Hayashi *et al*., 2012) revealed that rhizobia inoculation significantly enhanced the transcription of *GmTML1a/b* and *GmTML2* (Figure 4C and 4D). The increase in *GmTML1a/b* was seen following nodule emergence, corresponding to a developmental stage when AON would have taken effect (Figure 4C). It also coincides with the stage when miR2111 transported from the shoot would have decreased, enabling transcripts of TML to avoid degradation. Transcriptional analysis of wild-type roots from pot-grown plants revealed a similar regulatory trend for *TML* genes over a longer developmental period (Supplemental Figure 5).

### Target mimicry of *GmTML* genes and miR2111 overexpression alters nodule and root development

Target mimicry was employed to sequester mature miR2111 which would otherwise be available to target endogenous *GmTML* transcripts. Three short constructs, MIM2111_1, MIM2111_2 and MIM2111_3, were made (Figure 5A) and constitutively expressed in supernodulating *Gmnark* mutant hairy roots. Mimic MIM2111_1 was completely complementary with miR2111 and did not significantly alter nodule numbers compared to the empty vector (EV) control. This is consistent with what has been shown with mimics of other microRNAs (Todesco *et al*., 2010). In contrast, MIM2111_2 and MIM2111_3 did significantly reduce nodulation in *Gmnark* roots (Figure 5C). Cleavage of *GmTML1a*/*b* and *GmTML2* transcripts occurs between the 10^th^ and 11^th^ positions of miR2111 (Figure 5A) (Xu *et al*., 2013; Zhang *et al*., 2014), which is in agreement with MIM2111_2 being more effective at reducing nodulation than MIM2111_3 (Figure 5C). Nodules on empty vector-transformed control hairy roots of *Gmnark* composite plants were densely clustered and widely distributed throughout the roots (Figure 5F and 5J; the supernodulation phenotype), while MIM2111_2 formed considerably fewer nodules and lacked the stunted root phenotype of the mutant (Figure 5G; the autoregulated phenotype of wild-type plants). Constitutive MIM2111_2 expression also reduced nodulation in wild-type soybeans roots (Figure 5B).

**Figure 5.**
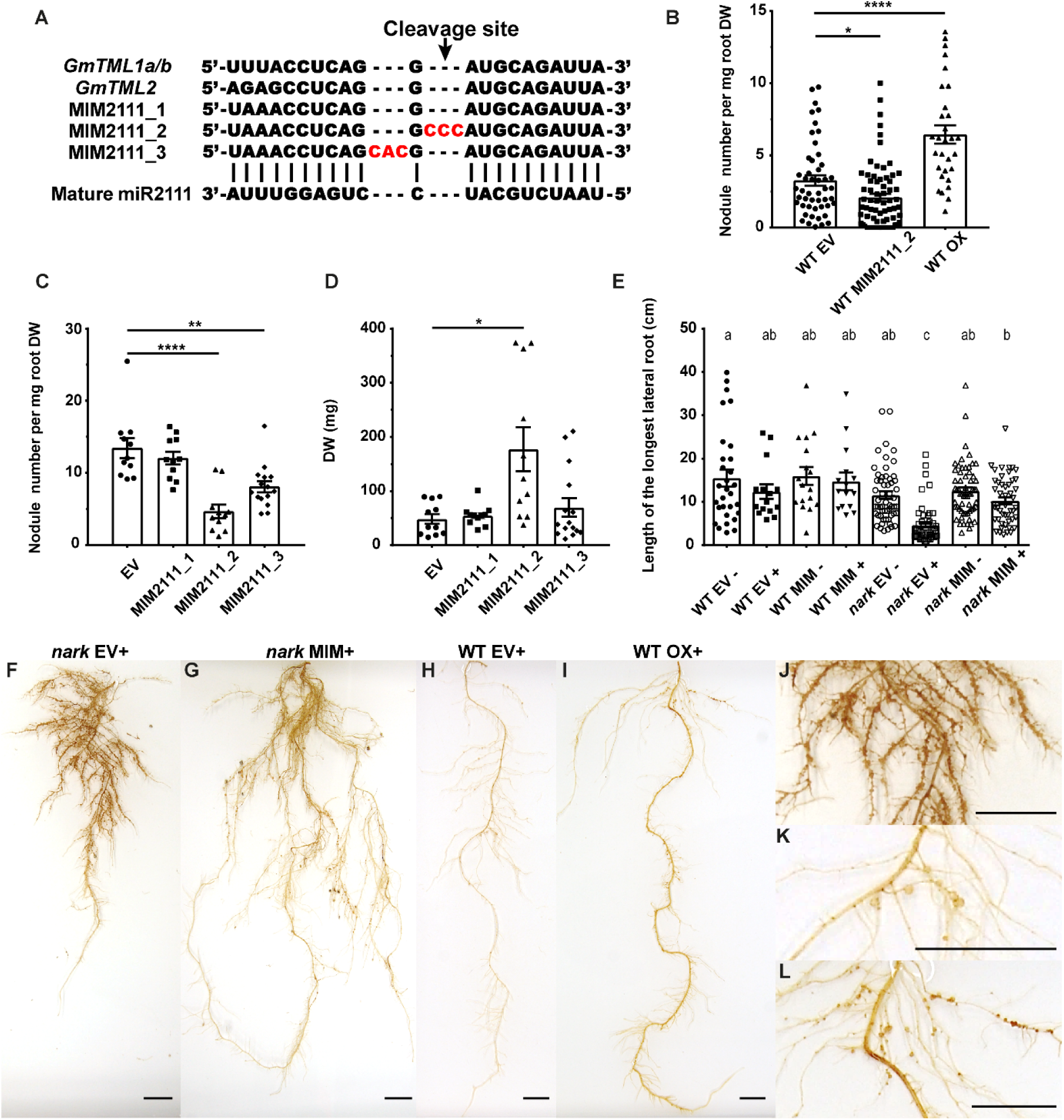
Nodule and root length phenotypes of transgenic supernodulating and wild-type soybean roots harbouring *GmTML* target mimic or miR2111 overexpression constructs. **(A)** Sequences of three target mimic constructs. MIM2111_1 is completely complementary to mature miR2111, while MIM2111_2 and 3 have a highlighted trinucleotide insertion (red) between the 10^th^ and 11^th^ positions, and the 11^th^ and 12^th^ positions of the miR2111 recognition sites of the *GmTML* transcripts, respectively. **(B)**Normalised nodule number of individual hairy roots transformed with EV, or MIM2111_2, or miR2111 overexpression (OX) construct in the wild-type background (*n* = 30-70). **(C and D)** Normalised nodule number **(C)** and root dry weight **(D)** of individual hairy roots of supernodulating *Gmnark* plants transformed with empty vector (EV), MIM2111_1, 2 or 3 (*n* = 11-15). **(E)** Length of the longest lateral root of individual EV and MIM2111_2 transformed hairy roots with (+) or without (-) *B. diazoefficiens* inoculation (*n* = 15-60). **(F-L)** Images of nodulating *Gmnark* or wild-type hairy roots transformed with EV, or MIM2111_2 or miR2111_OX construct. **J, K** and **L** are higher magnification images of **F, H** and **I** highlighting portions of roots exhibiting nodules, respectively. Scale bars = 2 cm. The experiments were repeated three times independently with similar results. Data are shown as mean ± SEM with different letters or asterisks indicating statistical significance. **(B-D)**: \**p* < 0.05, \*\**p* < 0.01, \*\*\*\**p* < 0.0001 (Student’s *t*-test). **(E)**: One-way ANOVA with post-hoc Tukey HSD test.

In addition to target mimicry of *GmTML* genes, ectopic overexpression of miR2111 was used to evaluate the microRNA’s function. Compared with EV control plants (Figure 5H and 5K), miR2111 over-expression strongly promoted nodulation in wild-type roots (Figure 5B, 5I and 5L). This is consistent with the target mimicry findings, and is congruent with miR2111 having a role in promoting nodulation by preventing the inhibitory mechanism of AON.

*Gmnark/Ljhar1/Mtsunn/Pssym29* mutants typically exhibit more lateral roots in the absence of rhizobia infection and severely stunted root growth following rhizobia inoculation (Day *et al*., 1986; Sagan and Duc, 1996; Wopereis *et al*., 2000; Schnabel *et al*., 2005). Indeed, these characteristics were also observed here (Figure 5F and 5J). However, the diminished root phenotype was significantly mitigated by constitutive MIM2111_2 expression (Figure 5G), with the dry weight of inoculated *Gmnark* hairy roots expressing MIM2111_2 significantly increased compared with EV control roots (Figure 5D). Likewise, the length of the longest lateral root was restored to that of the wild-type (Figure 5E). These evidence unequivocally demonstrated that miR2111 is the signalling factor acting downstream of GmNARK.

### Fine tuning the level of miR2111 regulates lateral root development

In wild-type plants, lowering miR2111 availability by target mimicry gave rise to slightly reduced lateral root density (*p* = 0.0368), while increased miR2111 expression stimulated lateral root emergence (*p* < 0.0001) (Figure 6A). Interestingly, first-order lateral roots of these miR2111 over-expressing hairy roots were considerably shorter than those of the EV control (Figure 6E). However, no statistically significant change in lateral root elongation was observed when MIM2111_2 was constitutively expressed (Figure 6E).

**Figure 6.**
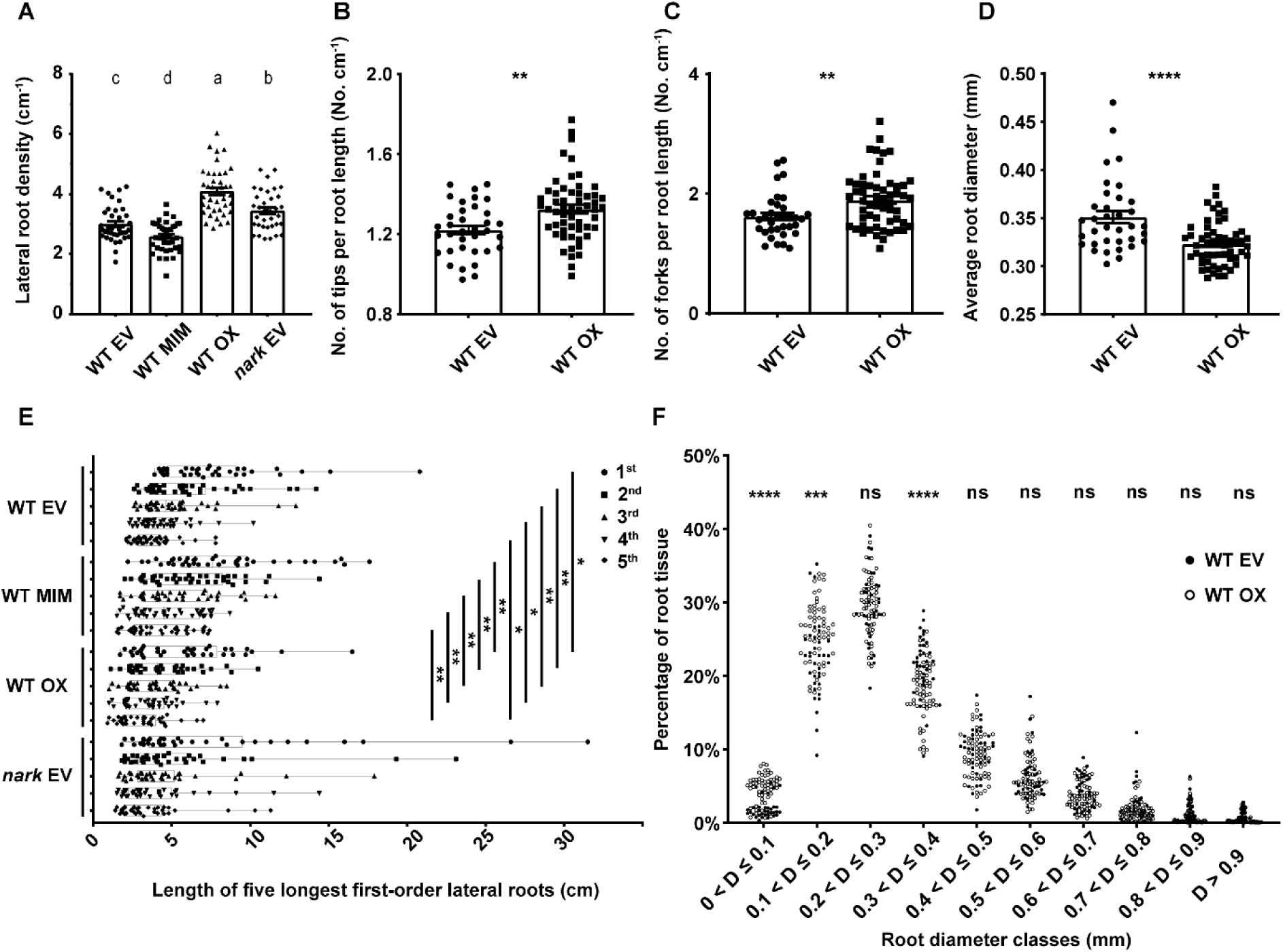
Root phenotyping of uninoculated supernodulating and wild-type soybean roots transformed with the EV, MIM2111_2 or miR2111_OX constructs. **(A)** Average number of emerged lateral roots per centimetre of root length in hairy roots transformed with EV, or MIM2111_2, or miR2111_OX construct and EV-expressing *Gmnark* hairy roots (*n* = 35-39). **(B and C)** Normalised number of root tips and forks (branching points) in miR2111_OX and EV-expressing hairy roots (*n* = 35-56). **(D)** Average root diameter of whole hairy roots expressing the EV or miR2111_OX construct (*n* = 35-56). **(E)** Length of five longest lateral roots in uninoculated hairy roots transformed with EV, or MIM2111_2, or miR2111_OX construct and EV-expressing *Gmnark* hairy roots. The borders of box plots indicate the lower and upper quartiles of the dataset with the lines inside the boxes and whiskers representing the medium and extrema (*n* = 35-39). **(F)** Percentage of roots falling into each root diameter class based on individual hairy roots transformed with EV or miR2111_OX construct (*n* = 35-56). The experiments were repeated three times independently with similar results. Data are shown as mean ± SEM with different letters or asterisks indicating statistical significance. **(A)**: One-way ANOVA with post-hoc Tukey HSD test. **(B-F)**: \**p* < 0.05, \*\**p* < 0.01, \*\*\**p* < 0.001, \*\*\*\**p* < 0.0001 (Student’s *t*-test). See also Supplemental Figure 6.

The ability of miR2111 to enhance root formation is demonstrated by a rise in the total number of root tips and forks (branching points) of hairy roots transformed to overexpress miR2111 (Figure 6B and 6C). This enhanced lateral root production was accompanied by a decrease in average root diameter (Figure 6D). Indeed, wild-type roots over-expressing miR2111 had a significantly larger proportion of thinner higher-order lateral roots (0 < Diameter ≤ 0.2 mm) than the EV control (Figure 6F). Additionally, the distance between root tips and the first emerged lateral roots was significantly shortened in miR2111 over-expressing roots of wild-type plants (Supplemental Figure 6). Taken together, miR2111 acts to regulate lateral root development, influencing the number of lateral roots, their thickness, the degree of root branching, and their positioning and emergence relative to the main root tip.

## Discussion

Identification of the systemic shoot-derived signal acting in AON to control legume nodule numbers has long been sought after. Here we use soybean to show miR2111 is the elusive signal that regulates both nodule organogenesis and root system architecture. The level of miR2111 is downregulated through GmNARK-dependent signalling in the leaf following rhizobia inoculation, leading to a reduction in mature miR2111 transported to the root. The reduced level of miR2111 in the root enables TML to accumulate, resulting in an inhibition of further nodule organogenesis. Collectively, these findings highlight that downregulation of leaf-derived miR2111, the expression of which was histologically co-localised with GmNARK (Nontachaiyapoom *et al*., 2007; Tsikou *et al*., 2018), is central to the AON pathway (Figure 7).

**Figure 7.**
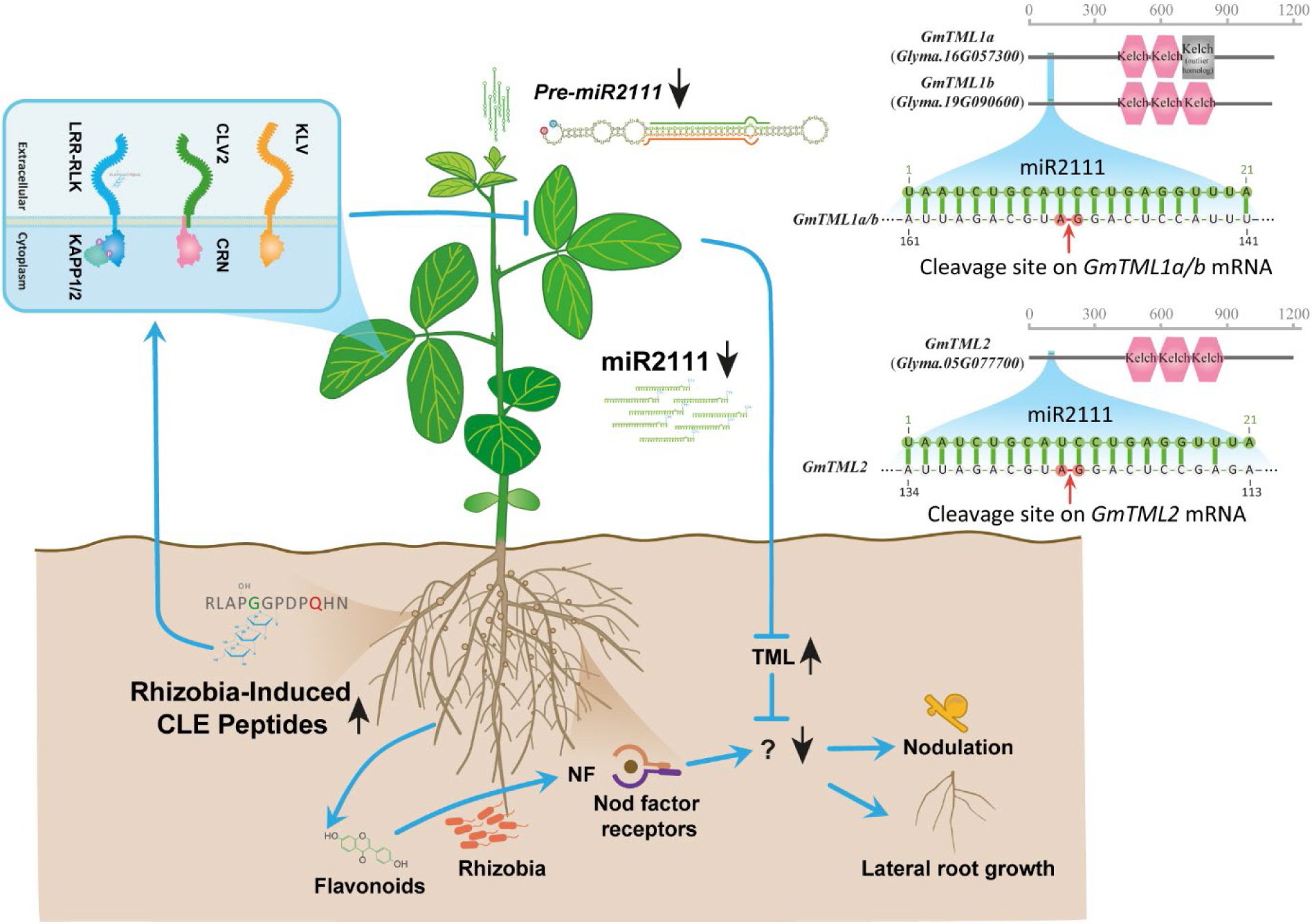
Proposed model of the AON pathway. Rhizobia infection triggers the production of nodulation-inhibitory CLE peptides in the root, which are transported via the xylem to the shoot where they are perceived by a LRR-RK (Sagan and Duc, 1996; Krusell *et al*., 2002; Searle *et al*., 2003; Schnabel *et al*., 2005; Ferguson *et al*., 2014), likely in a complex with other factors such as KLAVIER, CLAVATA2, and CORYNE (Miyazawa *et al*., 2010; Krusell *et al*., 2011; Crook *et al*., 2016). The kinase domain of the LRR-RK can be phosphorylated and de-phosphorylated by KAPP1 and 2, which likely have a role in downstream signal transduction (Miyahara *et al*., 2008). Perception of the CLE peptides by the LRR-RK downregulates the production of miR2111, which can be transported in the phloem to the roots where it mediates the degradation of the mRNA of Kelch repeat-containing F-Box proteins that inhibit nodulation. Example cleavage sites from soybean *GmTML1a/b* and *GmTML2* are shown. In AON, the reduced production of miR2111 in the shoot leads to an elevated level of the Kelch repeat-containing F-Box proteins in the root, resulting in the suppression of further nodule organogenesis. Arrows represent the regulatory change of each signal following the initiation of AON.

Although three *GmTML* genes have a pronounced transcriptional response to rhizobia infection, GmTML1 may have a predominant role over GmTML2 in the AON pathway when comparing their respective sequence identity with LjTML. This is further supported by the observation that only *GmTML1b* transcription was continuously elevated throughout nodule organogenesis. What factor(s) the TMLs target to suppress nodulation remains of interest to identify. To date, there is only one well-characterised family of Kelch repeat-containing F-Box proteins; these members contain an additional light-oxygen-voltage (LOV) domain and act to regulate protein stability in circadian clock and photoperiodic flowering pathways (Somers *et al*., 2000; Imaizumi *et al*., 2005; Ito *et al*., 2012).

*Ljtml-1 Ljhar1* double mutants exhibited no additive effect on the hyper-nodulation phenotype compared to the parental lines (Takahara *et al*., 2013), indicating that miR2111 is the sole shoot-derived factor functioning to restrict further nodule formation. This is consistent with our data where the nodule number and root length phenotypes of inoculated *Gmnark* mutant hairy roots transformed with MIM2111_2 were comparable with that of WT plants. Liberating *GmTML* transcripts from miR2111-mediated repression by over-expressing short target mimics of *GmTML* genes inhibited nodule development in wild-type and supernodulating mutant soybeans, further demonstrating that miR2111 acts through TML to regulate nodule numbers.

In contrast, overexpression of miR2111 resulted in a significant increase in nodule number. Similarly, petiole feeding synthetic double-stranded mature miR2111 promoted nodulation. Our findings are consistent with a recent study implicating a role for miR2111 in the AON pathway of *L. japonicus* (Tsikou *et al*., 2018) and support other studies demonstrating that systemically acting microRNAs are a common adaptive strategy for plants to cope with environmental stimuli (*e*.*g*., miR399 and miR395) (Pant *et al*., 2008; Buhtz *et al*., 2010; Marín-González and Suárez-López, 2012).

The miR2111 family is well conserved between many legume species, but it is not specific to legumes, suggesting its role in AON likely evolved from other developmental processes. In several model legume species, the precursor sequence of miR2111 has self-perpetuated by duplication, allowing for diversification and neofunctionalisation. Indeed, *Gm-miR2111* members appear to be preferentially expressed in either leaf or root tissue, and the genes within each group responded similarly to rhizobia inoculation. Our findings also indicate that the expression of miR2111-encoding genes correlates reasonably well with the level of mature miR2111.

A change in miR2111 abundance was associated with altered root development. Target mimicry of TML-encoding genes of soybean alleviated the aberrant growth of nodulating *Gmnark* mutant roots, and reduced the lateral root number of uninoculated wild-type roots. Moreover, constitutive miR2111 expression boosted lateral root emergence in wild-type plants, which contributed to the reduction in average root diameter and lateral root length. The co-regulation of lateral root and nodule formation by miR2111 suggests that this microRNA functions to promote the mitotic activity of meristematic cells in roots. In particular, the reduced spacing between lateral roots observed in miR2111 over-expressing roots reflects enhanced proliferation of lateral root founder cells. Additionally, these findings indicate that miR2111’s role in nodulation control may have evolved from a mechanism to regulate root system architecture in response to nutrient availability in the rhizosphere. This is consistent with the observations that miR2111 expression can respond to either phosphate or nitrate (Hsieh *et al*., 2009; Pant *et al*., 2009; Liang *et al*., 2012; Xu *et al*., 2013), two of the most influential factors on root system architecture (Péret *et al*., 2011; Forde *et al*., 2014). Therefore, recruiting miR2111 into the AON circuit of legumes may have evolved to connect nodulation control to the nutritional status of the host plant.

The discovery of miR2111 as the positive systemic regulator of nodule organogenesis has considerably bolstered the molecular framework of the AON pathway (Figure 7) (Ferguson *et al*., 2010). Future work to establish the protein target(s) of the TMLs is now of great interest, as it could help to elucidate the critical switch between achieving nodule development and nodulation control. Equally important is establishing the signalling relay between the LRR-RK of AON and the miR2111-encoding genes following perception of the nodulation-suppressive CLE peptides. As multiple transmembrane proteins are involved in ligand recognition and subsequent signal transduction events, it will be interesting to elucidate the contribution of individual components on the regulation of miR2111, which may assist in understanding the moderate downregulation of miR2111 observed in *Gmnark* leaves following rhizobia inoculation.

## Methods

### Plant materials

Wild-type (*Glycine max* [L.] Merr. cv. Bragg) and supernodulating *Gmnark* non-sense mutant (*nts382*) soybean seeds were chlorine gas sterilised for 16 hours and then germinated as outlined below for each experiment. For nodulation studies, plants were inoculated with *B. diazoefficiens* strain USDA110 or its non-nodulating isogenic mutant, *nodC*^*-*^ (Nieuwkoop *et al*., 1987), at OD600 ∼ 0.1.

### Bioinformatic analyses

BLASTN searches of all putative pre-miR2111 in *G. max Wm82*.*a2*.*v1, P. vulgaris v2*.*1, M. truncatula Mt4*.*0v1* and *A. thaliana TAIR10* genomes were conducted in Phytozome (https://phytozome.jgi.doe.gov/). This was initially performed using known pre-miR2111 members recorded in miRBase (http://www.mirbase.org/) and then subsequently repeated with precursor sequences identified here (E-value = 10). Likewise, *Lj*-miR2111 members were identified using the Legume Information System (LIS; https://legumeinfo.org/). As outlined previously (Hastwell *et al*., 2015; 2017), multiple sequence alignments of pre-miR2111 members were generated by Clustal Omega (http://www.ebi.ac.uk/Tools/msa/clustalo/), with slight manual adjustments mainly for the region corresponding to the conserved motif ∼150 bp upstream of the precursor sequences. The cladogram was constructed based on the above-mentioned alignment using 100 bootstrap support value as described previously (Hastwell *et al*., 2015; 2017). Genomic microsyntenies of *GmTML* gene members were achieved using the genomic context viewer of LIS which allows for the alignment of genes sharing similar microsyntenies. For miR2111 members, genomic environments were determined using the closest neighbouring genes found to be conserved across legumes. Manual matching was required when syntenies between three *GmTML* members or different miR2111 members could not be displayed in the same view.

### RNA extraction and quantitative real-time/stem-loop PCR

For microRNA quantification, two-week-old plants grown in 4 L pots containing sterile Grade 2 vermiculite provided with B&D nutrient solution (Broughton and Dilworth, 1971) supplemented with nitrogen (1 mM KNO3) were inoculated with *B. diazoefficiens* USDA110. The plants were grown for 2 more weeks, following which mature trifoliate leaves and roots from three individual plants grown in different pots were harvested and pooled for each biological replicate. The bottom 0.5 cm of the root was removed to exclude mRNA from the root tip which would dilute the sample. For *GmTML* transcriptional analysis, three-day-old seedlings were transferred to modified CYGTM germination pouches, with the pouch length extended to prevent roots from reaching the bottom (Mega International, Newport, MN, USA) (Corcilius *et al*., 2017) and acclimatised for two days in a TPG-1260-TH Thermoline growth chamber (16-hour photoperiod and 28°C:24°C day/night), as described previously (Hayashi *et al*., 2012). 1 mL of *B. diazoefficiens* USDA110 culture was slowly applied to the Zone Of Nodulation (ZON) of the tap root, with the location of the root tip marked at the time of inoculation. Water was provided to the pouch twice a day as required to prevent drying of the pouch filter paper. A 5 cm region of the tap root around the marked site was harvested from three individual plants and pooled for each biological replicate. For leaf and root samples, total RNA was extracted using Maxwell® RSC Plant RNA or Maxwell® 16 miRNA Tissue Kit from Promega according to the manufacturer’s instructions. Diluted cDNA synthesized from 1 µg of RNA (Invitrogen™; SuperScript™ III First-Strand Synthesis System) was used for qRT-PCR with *GmCons6* used as the reference housekeeping gene (Libault *et al*., 2008; Udvardi *et al*., 2008). Mature miR2111 and miR1520d were reverse transcribed using long stem-loop reverse primers from an equal amount of RNA as described previously (Varkonyi-Gasic *et al*., 2007) and stem-loop RT-PCR was performed with gene-specific forward and universal reverse primers with miR1520d as the housekeeping microRNA (Kulcheski *et al*., 2010). Absolute stem-loop RT-PCR was used to quantify mature miR2111 in soybean phloem sap due to a lack of reference microRNAs reported for this sample type. Primer sequences for quantitative PCR are listed in Supplemental Table 2. Initially, 800 µL of sap was centrifuged at 4°C for 3 min to eliminate tissue debris before transfer to a new tube and then mixed with 300 µL of Lysis buffer of the Maxwell® RSC Plant RNA Kit. The entire 1.1 mL was used for RNA extraction.

### Construct creation

Target mimic constructs were made by directly ligating the oligo duplexes generated from three complementary primer pairs (Supplemental Table 2) to the *Eco*RI and *Hin*dIII double-digested integrative vector (p15SOG; unpublished). The ligation products were transformed into XL1-BLUE competent *E. coli* cells using heat shock and then transformed cells were cultured overnight on LB plates containing 100 µg/ml spectinomycin. Colony PCR was performed to identify putative positive clones from which plasmids were extracted for further confirmation via sequencing. An additional construct containing the full amplicon sequence was created for absolute stem-loop RT-PCR of mature miR2111. The annealed primer duplex (Supplemental Table 2) was ligated into the linearized pGEM-T^®^ Easy vector. The ligated product was transformed into XL1-BLUE competent *E. coli* cells. Plasmids extracted from positive bacterial clones confirmed by colony PCR and sequencing were used to make dilution series. To generate the construct used for *in-vivo* miR2111 synthesis, a pair of long complementary primers (Supplemental Table 2) which can form a perfectly matched stem-loop structure corresponding to an artificial miR2111 precursor was designed. These primers include sticky ends needed for subsequent ligation with digested vectors and could be forced to anneal by gradually lowering the temperature from 98°C. Modification of the vector, L4440 (Mitter *et al*., 2017), was made to replace one of the T7 promoters with a T7 terminator. In addition, the sequence previously flanked by two T7 promoters was replaced with two adjacent *Bsa*I restriction sites. This new vector generated here is named L4440T7ter. Ligation of *Bsa*I digested L4440T7ter with the primer duplex enables considerable production of the artificial pre-miR2111 under the control of the IPTG-inducible T7 promoter. The ligation product was transformed into the HT115(DE3) bacterial strain using heat shock and positive clones were selected by liquid colony PCR and confirmed by sequencing. For the miR2111 overexpression construct, the precursor sequence of *Gm-miR2111a* was PCR amplified using the primers listed in Supplemental Table 2. Following column purification, the *Eco*RI and *Hin*dIII double-digested PCR fragments and p15SOG vector were ligated and subsequently transformed into XL1-BLUE competent *E. coli* cells for further identification as described above.

### Triparental mating

Following PCR confirmation, auxotrophic HB101 *E. coil* transformed with the recombinant plasmids (spectinomycin resistance), helper plasmid pRK2013 (kanamycin resistance) containing *E. coli* strain, and *Agrobacterium rhizogenes* strain K599 (rifampicin resistant) were cultured separately in liquid LB medium with their respective appropriate antibiotic overnight. Bacterial pellets obtained from 1 mL of cell culture centrifuged at 13,000 rpm (Eppendorf 5424 Centrifuge with Fa-45-11 Rotor) were thoroughly washed three times with 1 mL fresh LB medium to remove residual antibiotics and were re-suspended in 100 µL liquid LB medium. Three bacterial strains were mixed at a ratio of 1:1:1 and subsequently pipetted onto plain LB plates for triparental mating. Following one-day incubation at 28°C, the bacterial mixture was re-streaked onto minimal (MIN) medium containing 100 µg/ml spectinomycin and 50 µg/ml rifampicin. To reduce the background growth of unintended bacterial strains, single colonies from MIN plates incubated at 28°C for two days were sub-cultured overnight in 1 mL liquid MIN medium having the same concentration of both antibiotics and the bacteria grown in liquid culture were subsequently streaked onto antibiotic-containing MIN plates. Positive clones tested by PCR were selected for hairy root transformation.

### Hairy-root transformation

Soybean seeds were sown in pots half filled with Grade 2 vermiculite at 28°C:24°C in a TPG-1260-TH Thermoline growth chamber. Four-day-old seedlings were stabbed using a fine needle filled with thick *A. rhizogenes* suspension just beneath the unopened cotyledons (Kereszt *et al*., 2007) and transferred to 25°C:22°C day/night temperature to encourage hairy-root growth. The pots were covered with transparent cellophane wrap to induce high humidity and after three days additional vermiculite was added to bury the wound site. Every three days, B&D nutrient solution supplemented with 1 mM KNO3 was supplied to the plants. After three weeks, the original non-transgenic root system was excised and the remaining plant with emerging hairy roots was transplanted into new pots at 28°C:24°C day/night with a 16-hour photoperiod. Three days later, the plants were inoculated with *B. diazoefficiens*. For Figure 5B-5D, nodule numbers were scored two week-post-inoculation, whilst other data in Figure 5 were obtained from hairy roots harvested three week-post-inoculation. Root phenotyping occurred at two (Figure 6A and 6E) and 3 weeks (Figure 6B-6D, and 6F; Supplemental Figure 6) post transplantation. Data presented in Figure 6B, 6C, 6D and 6F were determined using WinRhizo (version: Arabidopsis 2019a).

### *In-vivo* miRNA synthesis

To generate microRNA, HT115(DE3) bacteria harbouring recombinant L4440T7ter plasmids for artificial pre-miR2111 were grown overnight in 200 µg/ml ampicillin and 12.5 µg/ml tetracycline-containing liquid LB medium and used for 1:100 inoculation of 2xYT medium (100 µg/ml ampicillin and 12.5 µg/ml tetracycline). The bacteria were grown at 37°C in shakers to OD600 = 0.4-0.5, at which time IPTG was added to the cultures. After a further two-hours of incubation with 0.5 mM IPTG, the total concentration of IPTG in the culture was increased to 1 mM, followed by an additional two-hour incubation to allow for the induction of gene expression. The bacteria were subsequently pelleted by centrifugation (Sigma 4K15 Centrifuge), the medium discarded, and the pellet stored at -20°C. The bacteria were re-suspended in a small amount of Milli-Q® water, mixed with an equal amount of phenol:chloroform:isoamyl alcohol (25:24:1; pH6.7/6.8), vortexed vigorously and left to stand for 0.5-1 hour at room temperature. The mixture was centrifuged (Eppendorf 5424 Centrifuge) at >13,000 rpm for at least ten minutes to ensure complete phase partitioning. The upper clear aqueous phase was transferred to a new tube and thoroughly mixed with an equal amount of isopropanol. Precipitation occurred overnight at -20°C and the RNA pellet was obtained by centrifugation at 3,000 rpm at 4°C for ten minutes (Sigma 4K15 Centrifuge). After discarding the supernatant, RNA samples were washed using 75% ethanol and pelleted by centrifugation with increased speed to remove ethanol. This washing step was repeated three times. The pellet was then heated at 65°C to fully evaporate remaining ethanol, after which it was dissolved in small amount of RNase-free water on ice. RNA extracted from approximately 250 mL of bacterial culture grown in 37°C shakers was then treated with 10 µL of DNase I (Fermentas). After one-hour of digestion, the total volume of the sample was doubled using 1 M NaCl before addition of 10 µL of RNase A (Fermentas), thereby specifically preventing digestion of double-stranded RNA under high salt condition as opposed to single-stranded RNA. Following one-hour incubation at 37°C, the processed RNA samples were mixed with an equal amount of phenol:chloroform:isoamyl alcohol (25:24:1; pH6.7/6.8) and subsequently purified as described above.

### Petiole feeding

For petiole feeding synthetic double-stranded miR2111, wild-type soybean plants were grown at 28°C:24°C day/night and 16 h:8 h day/night in an E-75L1 Percival growth cabinet. Three-week-old plants having two mature trifoliate leaves were fed using a petiole feeding apparatuses as described previously (Lin *et al*., 2010; 2011; Hastwell *et al*., 2019). The volume of solution taken up by each recipient plant was recorded. RNA samples being fed through the syringes were replenished each day and refreshed every three days. The recipient plants were inoculated with *B. diazoefficiens* USDA110 after being fed for one day. Throughout the experiment, plants were watered every three days with B&D nutrient solution supplemented with 2 mM nitrate to promote growth without inhibiting nodulation. As the root systems were already well developed in medium-sized pots prior to miRNA feeding, no statistically significant root architectural differences were detected between the treatments and controls at the time of harvest.

### RNA sequencing

For RNAseq, the first trifoliate leaves of wild-type soybean plants were harvested one-week after inoculation with either *B. diazoefficiens* USDA110 or its *nodC*^*-*^ mutant. PureLink™ DNase-treated RNA samples were isolated from ground leaf tissue using the Invitrogen™ PureLink™ RNA Mini Kit and used to make cDNA libraries according to the Illumina TruSeq^®^ Stranded mRNA Library Preparation Kit. Samples prepared from two biological replicates for each inoculation type were sequenced via Illumina HiSeq2500. The paired-end reads were mapped using Geneious v11.0.2 with a customized sensitivity setting (Minimum mapping quality: 30; Word length: 18; Index word length: 13; Maximum mismatches per read: 1%; Maximum ambiguity: 4). Sequencing data that was generated in this study have been deposited in Gene Expression Omnibus (GEO) with the accession codes PRJNA591309 and SAMN13379533.

### Sap collection

For sap collection, two-week old soybean plants were inoculated with *B. diazoefficiens* USDA110 and subsequently harvested at 14 dpi. Sap was extracted from stem segments of inoculated and uninoculated soybean plants using centrifugation. Multiple stem segments (∼2 cm) were placed vertically in 0.6 mL tubes, which were then centrifuged for 3 min at 4°C. The sap collected was transferred to 2 mL RNase-free Eppendorf tubes and stored at -80°C.

### DNA extraction

A small quantity of ground root tissue was lysed using 65 - 80 µL of TES buffer (100 mM Tri-HCl (pH = 8.0), 2 mM EDTA, 20 g/L SDS) and an equal volume of acidic phenol:chloroform:isoamyl alcohol (25:24:1). Following 10-minute incubation at 70°C, the mixture was centrifuged at 12,500 rpm for five minutes and 40 µL of the top clear aqueous phase was immediately added onto the Sephadex G-50 columns made in house. Trace amounts of DNA were obtained after five-minute centrifugation at 6,000 rpm and used for PCR confirmation of transgenic hairy roots.

### PAGE gel electrophoresis

To confirm the production of double-stranded miR2111 after serial enzymatic treatments, 12% non-denaturing PAGE gel consisting of 4.8 mL of 30% Acrylamide:Bis (29:1), 3.6 mL of 80% glycerol, 2.4 mL of 5X TBE buffer, 208 µL of 10% APS, 15 µL of TEMED was used. The gels were merged into the fixing solution (100 mL ethanol and 5 mL acetic acid/L) for 10 mins and subsequently stained in 0.2% AgNO3 solution for 10 mins in the dark. After rinsing with distilled water for a few times, the gel was transferred to a plastic plate containing 50 mL of the developer solution (7.5 g NaOH and 0.6 g Na2B4O7•10H2O/L) into which 150 µL of 37% formaldehyde was added to initiate imaging.

### Statistical analysis

All analyses were conducted using GraphPad Prism version 7.03. Statistical significance regarding transcriptional variation of *Gm-miR2111* and *GmTML* gene members, nodule number examination of petiole-fed plants and root phenotypic changes of target mimic/overexpression construct transformed soybeans was determined using one-way ANOVA coupled with post-hoc Tukey HSD test. Two-tailed Student’s *t*-test was performed to assess the expression analysis of mature miR2111, and the observed nodule number and root phenotypic differences between hairy roots transformed with EV, target mimic, overexpression constructs. All data are presented as mean ± SEM. The number of biological replicates used in each experiment is indicated in the figures as individual data points and/or reported as n in the figure legends.

## Author contributions

M.Z., P.M.G. and B.J.F. designed the project. M.Z. and H.S. performed the experiments. M.Z., H.S. and B.J.F. analysed the data. M.Z. and B.J.F. wrote the manuscript. H.S. analysed the RNAseq data and made Figure 7. All authors helped edit the manuscript.

### Acknowledgements

We thank Xitong Chu, Dongxue Li, April H. Hastwell, Audrey McInnerney, Leslie Macas, Zian Zhan, Celine Mens, Candice Jones for technical assistance and advice. Thanks also to Prof. Neena Mitter for the provision of the L4440 vector and the HT115 bacterial strain.

