## Supplemental Figure 1-6 & Table 1-3 for "Shoot-derived miR2111 controls legume root and nodule development"

### Supplemental information

#### Supplemental Figure 1

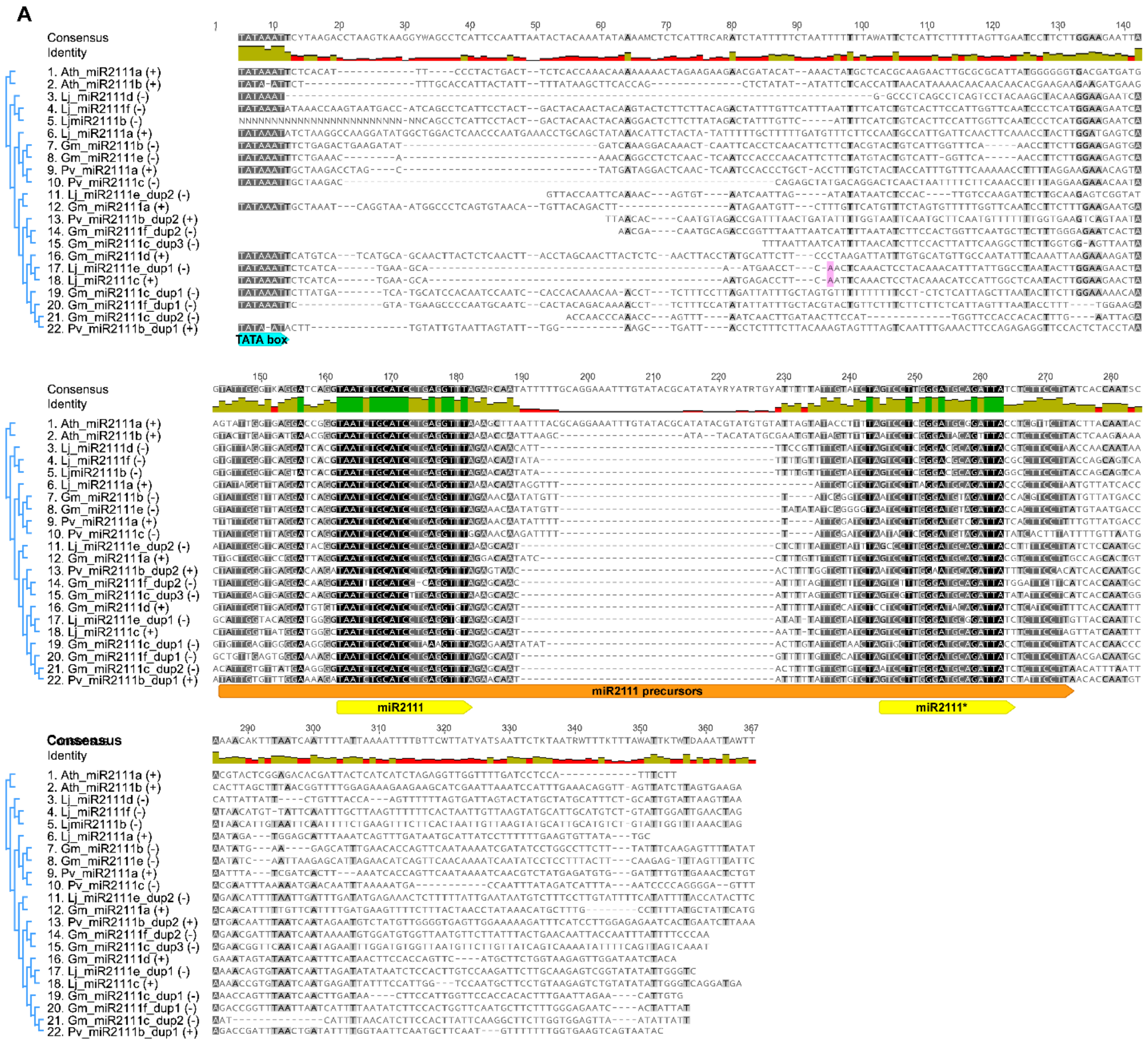

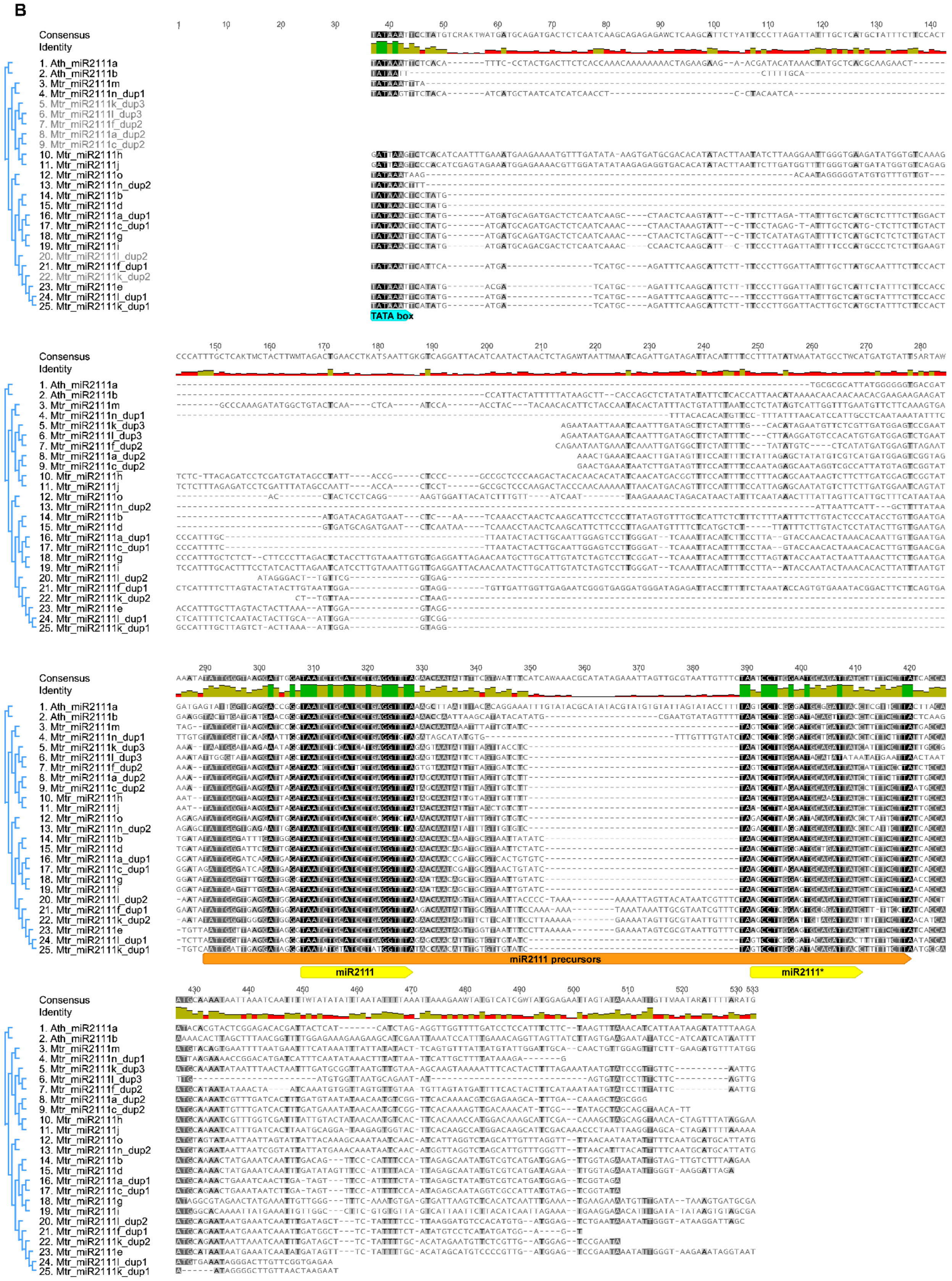
B
Consensus
Identity
1. Ath_miR2111a
2. Ath_miR2111b
3. Mtr_miR2111m
4. Mtr_miR2111n_dup1
5. Mtr_miR2111k_dup3
6. Mtr_miR2111l_dup3
7. Mtr_miR2111f_dup2
8. Mtr_miR2111a_dup2
9. Mtr_miR2111c_dup2
10. Mtr_miR2111h
11. Mtr_miR2111j
12. Mtr_miR2111o
13. Mtr_miR2111n_dup2
14. Mtr_miR2111b
15. Mtr_miR2111d
16. Mtr_miR2111a_dup1
17. Mtr_miR2111c_dup1
18. Mtr_miR2111g
19. Mtr_miR2111i
20. Mtr_miR2111l_dup2
21. Mtr_miR2111f_dup1
22. Mtr_miR2111k_dup2
23. Mtr_miR2111e
24. Mtr_miR2111l_dup1
25. Mtr_miR2111k_dup1
TATA box
miR2111 precursors
miR2111
miR2111*

**Supplemental Figure 1. Multiple sequence alignments of pre-miR2111 members of *A. thaliana* with either (A) *G. max*, *P. vulgaris*, and *L. japonicus* or (B) *M. truncatula*.**

The miR2111 precursors are shown in orange and sequences corresponding to mature miR2111 and miR2111* are shown in yellow. Cyan arrowheads indicate the position of the predicted TATA box. The specific transcription start sites of two *Lj-miR2111* genes confirmed by 5' RACE (Tsikou et al., 2018) are highlighted in pink.

**Supplemental Figure 2**

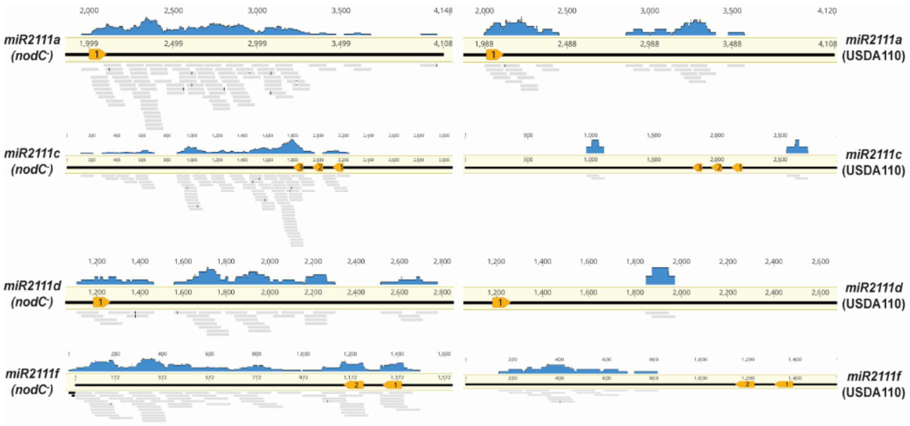


**Supplemental Figure 2. *Gm-miR2111* RNAseq read mapping of leaves harvested from soybean plants inoculated with either USDA110 or *$nodC^-$* rhizobia.**

*G. max* genomic sequences containing miR2111-encoding genes are indicated as a black line with orange arrowheads representing different miR2111 precursors. *Gm-miR2111b* and *e* were absent as almost no reads were detected, consistent with the RT-PCR analysis in Figure 2. Pooled from two biological replicates consisting of eight plants each, individual RNAseq reads shown as grey lines were mapped to the genomic sequence using very high stringency (≤ 1 mismatch).

**Supplemental Figure 3**

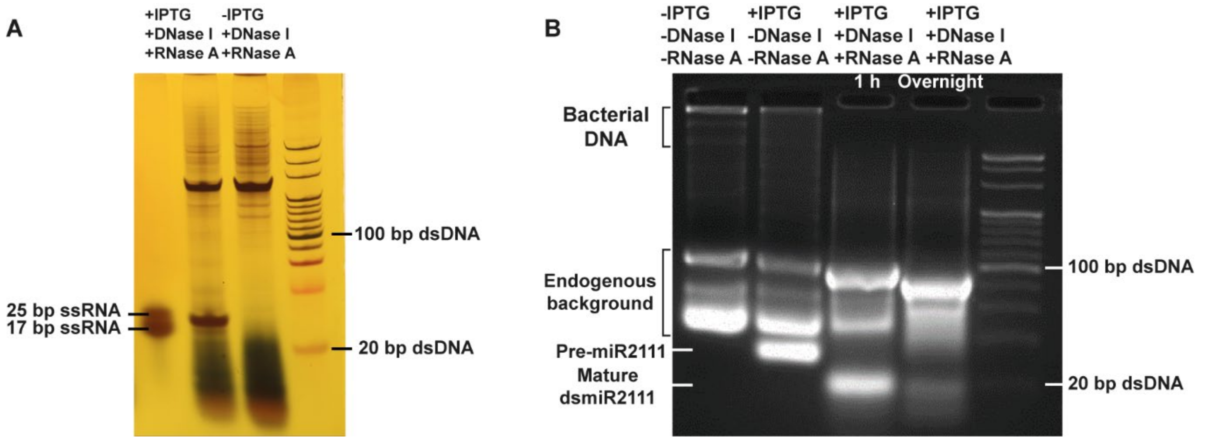


**Supplemental Figure 3. Gel photographs of artificial pre-miR2111 and double-stranded miR2111 produced *in vivo*.**

**(A)** Induced (+IPTG) and non-induced (-IPTG) enzymatically digested RNA samples on non-denaturing PAGE gel. One single-stranded RNA and a double-stranded DNA markers were included to indicate the size of the double-stranded miR2111 product.

**(B)** Non-induced and induced RNA samples with (+) or without (-) DNase I and RNase A treatment were run on 4% agarose gel along with a 20 bp marker. Samples in the third and fourth lanes were treated with DNase I for one hour. These samples were then subjected to RNase A treatment for one hour (lane 3) or overnight (lane 4).

**Supplemental Figure 4**

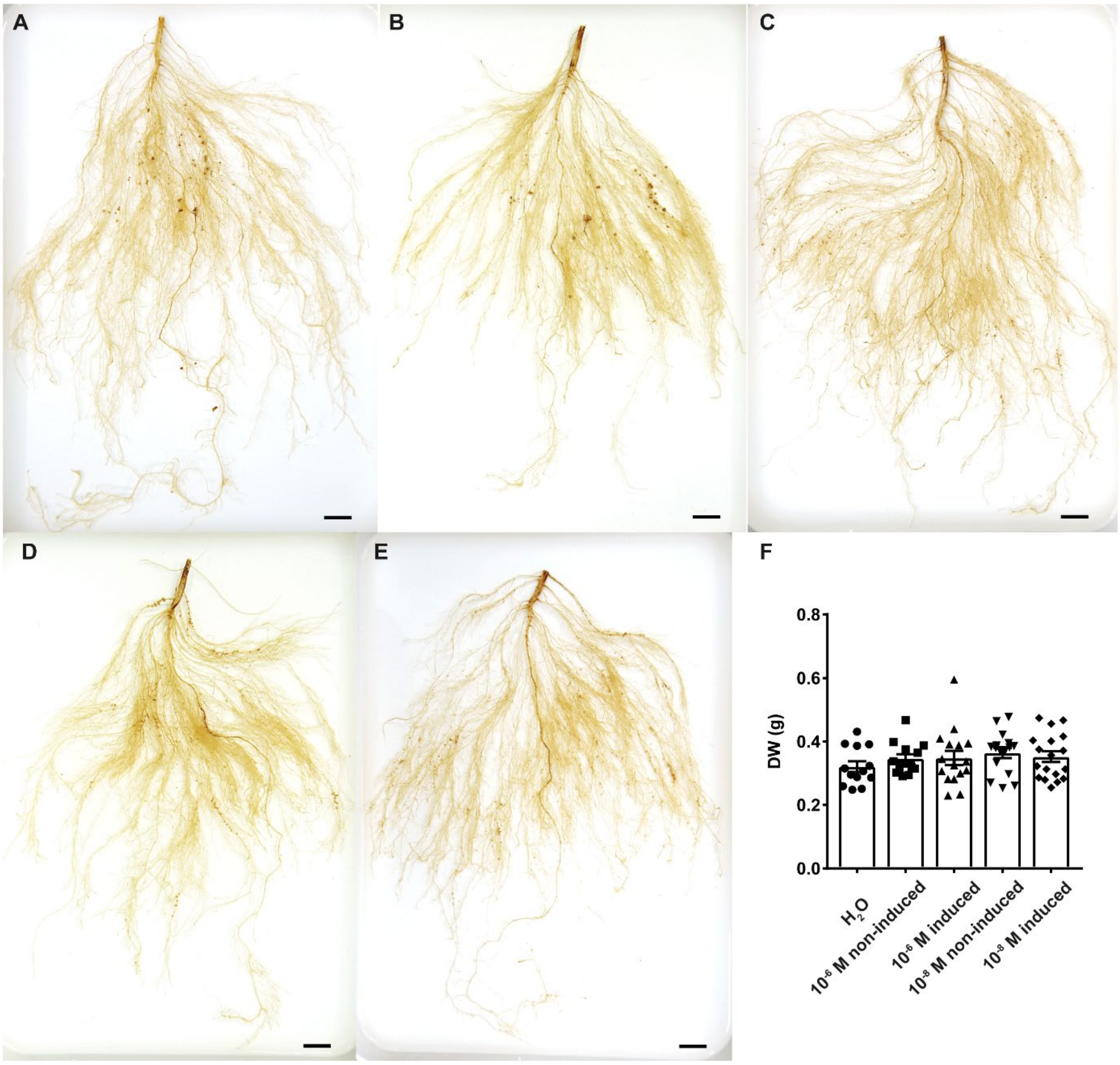


**Supplemental Figure 4. Images of a representative whole root system and root dry weight measurement from each treatment of the petiole feeding experiment, related to Figure 3.** **(A)** $H_2O$ **(B)** $10^{-6}$ M non-induced **(C)** $10^{-6}$ M induced **(D)** $10^{-8}$ M non-induced **(E)**$10^{-8}$ M induced. Scale bars represent 2 cm.

**(F)** Average whole root dry weight of the petiole-fed wild-type soybean plants. Data are shown as mean ± SEM (One-way ANOVA with post-hoc Tukey HSD test).

**Supplemental Figure 5**

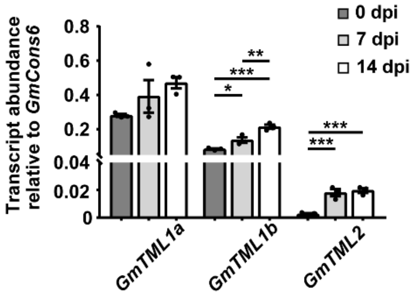


**Supplemental Figure 5. Transcriptional responses of *GmTML* family members during nodule development, related to Figure 4.**

Data shown are mean ± SEM ($n$ = 3 where each biological replicate was pooled from three wild-type plants grown in different pots. Asterisks denote statistically significant differences (*$p < 0.05$, **$p < 0.01$, ***$p \leq 0.001$; One-way ANOVA with post-hoc Tukey HSD test).

**Supplemental Figure 6**

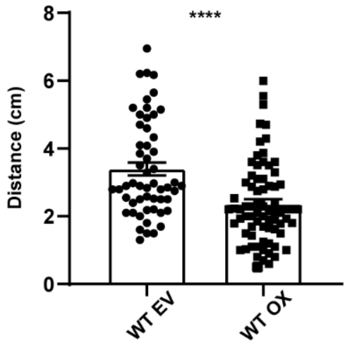


**Supplemental Figure 6. Proximity of root tips and the position of first emerged lateral roots from the parental roots in miR2111 over-expressing hairy roots.** Data are shown as mean ± SEM with asterisks indicating statistical significance ($n$ = 55-74, ****$p$ < 0.0001, Student's $t$-test).

**Supplemental Table 1**

**Genomic location and orientation of pre-miR2111 members identified in *L. japonicus* and *M. truncatula*.**

| Name of miR2111 precursors | Genomic location | Orientation on the chromosome |
|---|---|---|
| **Lj_miR2111a** | Lj1: 39791749 - 39791831 | (+) |
| **Lj_miR2111b** | Lj0: 77855203 - 77855118 | (-) |
| **Lj_miR2111c** | Lj0: 109196808 - 109196888 | (+) |
| **Lj_miR2111d** | Lj0: 129665012 - 129664927 | (-) |
| **Lj_miR2111e_dup1** | Lj0: 129674804 - 129674723 | (-) |
| **Lj_miR2111e_dup2** | Lj0: 129674638 - 129674556 | (-) |
| **Lj_miR2111f** | Lj0: 162789315 - 162789230 | (-) |
| **Mtr_miR2111a_dup1** | Mtr7: 23625581 - 23625482 | (-) |
| **Mtr_miR2111a_dup2** | Mtr7: 23625404 - 23625307 | (-) |
| **Mtr_miR2111b** | Mtr7: 23619774 - 23619674 | (-) |
| **Mtr_miR2111c_dup1** | Mtr7: 23614814 - 23614705 | (-) |
| **Mtr_miR2111c_dup2** | Mtr7: 23614628 - 23614531 | (-) |
| **Mtr_miR2111d** | Mtr7: 23607487 - 23607387 | (-) |
| **Mtr_miR2111e** | Mtr7: 23600218 - 23600081 | (-) |
| **Mtr_miR2111f_dup1** | Mtr7: 23590077 - 23589942 | (-) |
| **Mtr_miR2111f_dup2** | Mtr7: 23589870 - 23589773 | (-) |
| **Mtr_miR2111g** | Mtr7: 23584827 – 23584718 | (-) |
| **Mtr_miR2111h** | Mtr7: 23584441 – 23584344 | (-) |
| **Mtr_miR2111i** | Mtr7: 23578583 – 23578474 | (-) |
| **Mtr_miR2111j** | Mtr7: 23577905 – 23577809 | (-) |
| **Mtr_miR2111k_dup1** | Mtr7: 23573110 – 23573011 | (-) |
| **Mtr_miR2111k_dup2** | Mtr7: 23572987 – 23572850 | (-) |
| **Mtr_miR2111k_dup3** | Mtr7: 23572783 – 23572677 | (-) |
| **Mtr_miR2111l_dup1** | Mtr7: 23566411 – 23566305 | (-) |
| **Mtr_miR2111l_dup2** | Mtr7: 23566282 – 23566146 | (-) |
| **Mtr_miR2111l_dup3** | Mtr7: 23566077 – 23565971 | (-) |
| **Mtr_miR2111m** | Mtr7: 23550197 – 23550091 | (-) |
| **Mtr_miR2111n_dup1** | Mtr4: 6069052 - 6068942 | (-) |
| **Mtr_miR2111n_dup2** | Mtr4: 6068876 - 6068779 | (-) |
| **Mtr_miR2111o** | Mtr4: 6073342 - 6073245 | (-) |

**Supplemental Table 2**

**Statistical data for Figure 6.**

| Figure 6A | | | | |
|---|---|---|---|---|
| | WT EV | WT MIM | WT OX | *nark* EV |
| WT EV | - | - | - | - |
| WT MIM | Adjusted *P* = 0.0368 | - | - | - |
| WT OX | Adjusted *P* < 0.0001 | Adjusted *P* < 0.0001 | - | - |
| *nark* EV | Adjusted *P* = 0.0083 | Adjusted *P* < 0.0001 | Adjusted *P* = 0.0001 | - |

| Figure 6E | | | | | |
|---|---|---|---|---|---|
| | WT EV 1$^{st}$ | WT EV 2$^{nd}$ | WT EV 3$^{rd}$ | WT EV 4$^{th}$ | WT EV 5$^{th}$ |
| WT MIM 1$^{st}$ | *P* = 0.3905 | - | - | - | - |
| WT MIM 2$^{nd}$ | - | *P* = 0.7429 | - | - | - |
| WT MIM 3$^{rd}$ | - | - | *P* = 0.5541 | - | - |
| WT MIM 4$^{th}$ | - | - | - | *P* = 0.3797 | - |
| WT MIM 5$^{th}$ | - | - | - | - | *P* = 0.3069 |
| WT OX 1$^{st}$ | *P* = 0.0321 | - | - | - | - |
| WT OX 2$^{nd}$ | - | *P* = 0.0070 | - | - | - |
| WT OX 3$^{rd}$ | - | - | *P* = 0.0091 | - | - |
| WT OX 4$^{th}$ | - | - | - | *P* = 0.0183 | - |
| WT OX 5$^{th}$ | - | - | - | - | *P* = 0.0395 |
| *nark* EV 1$^{st}$ | *P* = 0.9429 | - | - | - | - |
| *nark* EV 2$^{nd}$ | - | *P* = 0.4247 | - | - | - |
| *nark* EV 3$^{rd}$ | - | - | *P* = 0.4021 | - | - |
| *nark* EV 4$^{th}$ | - | - | - | *P* = 0.6782 | - |
| *nark* EV 5$^{th}$ | - | - | - | - | *P* = 0.6915 |
| | WT MIM 1$^{st}$ | WT MIM 2$^{nd}$ | WT MIM 3$^{rd}$ | WT MIM 4$^{th}$ | WT MIM 5$^{th}$ |
| WT OX 1$^{st}$ | *P* = 0.0041 | - | - | - | - |
| WT OX 2$^{nd}$ | - | *P* = 0.0019 | - | - | - |
| WT OX 3$^{rd}$ | - | - | *P* = 0.0029 | - | - |
| WT OX 4$^{th}$ | - | - | - | *P* = 0.0035 | - |
| WT OX 5$^{th}$ | - | - | - | - | *P* = 0.0062 |

| Figure 6F | | | | | | | | | |
|---|---|---|---|---|---|---|---|---|---|
| 0<D≤0.1 | 0.1<D≤0.2 | 0.2<D≤0.3 | 0.3<D≤0.4 | 0.4<D≤0.5 | 0.5<D≤0.6 | 0.6<D≤0.7 | 0.7<D≤0.8 | 0.8<D≤0.9 | D>0.9 |
| *P* < 0.0001 | *P* = 0.0008 | *P* = 0.6280 | *P* < 0.0001 | *P* = 0.2892 | *P* = 0.5467 | *P* = 0.1360 | *P* = 0.0727 | *P* = 0.1653 | *P* = 0.1097 |

#### Supplemental Table 2

#### List of primers.

| Name of the primers | Sequence (5’ → 3’) | Use |
|---|---|---|
| **MIM2111_1 Fw** | AATTCTAAACCTCAGGATGCAGATTAG | Mimic construct creation |
| **MIM2111_1 Rv** | GATCCTAATCTGCATCCTGAGGTTTAG | Mimic construct creation |
| **MIM2111_2 Fw** | AATTCTAAACCTCAGGCCCATGCAGATTAG | Mimic construct creation |
| **MIM2111_2 Rv** | GATCCTAATCTGCATGGGCCTGAGGTTTAG | Mimic construct creation |
| **MIM2111_3 Fw** | AATTCTAAACCTCAGCACGATGCAGATTAG | Mimic construct creation |
| **MIM2111_3 Rv** | GATCCTAATCTGCATCGTGCTGAGGTTTAG | Mimic construct creation |
| **p15SOG_35s_Fw** | GAGGAGCATCGTGGAAAAAG | Colony PCR |
| **p15SOG_Rv** | GGCGGTAAGGATCTGAGCTA | Colony PCR/sequencing |
| **GmCons6_Fw** | AGATAGGGAAATGGTGCAGGT | qPCR |
| **GmCons6_Rv** | CTAATGGCAATTGCAGCTCTC | qPCR |
| **GmTML1a/1b_Fw** | CTAATGGAGATGAACCTTTACCTCAG | qPCR |
| **GmTML1a_Rv** | TAGATCTCGCCACTCCTCGCA | qPCR |
| **GmTML1b_Rv** | TAGATCTCGCCACTCCTTGAG | qPCR |
| **GmTML2_Fw** | AGTCACCTTCATATGCCCATGG | qPCR |
| **GmTML2_Rv** | GTCAGAAACCGCTTGCTTAGAC | qPCR |
| **GmMIR2111a_Fw** | CTTTGAAGAATGATTGCTGGGTCC | qPCR |
| **GmMIR2111a_Rv** | AGTGCTGGCAAGGAAGACG | qPCR |
| **GmMIR2111b/e_Fw** | GAAGAGTGAGTATTGGTTTAGGATCAG | qPCR |
| **GmMIR2111b_Rv** | GCTCTTCATATTGGTCATAACATAAGGAC | qPCR |
| **GmMIR2111c_Fw** | GGTGGAGTTAATATTATTGAGTGAGGACA | qPCR |
| **GmMIR2111c_Rv** | CACATCCAAATTCTATTGATTGAACCG | qPCR |
| **GmMIR2111d_Fw** | TGCATCCTGAGGTGTAGAGCA | qPCR |
| **GmMIR2111d_Rv** | CAACTCTTACCAGAAGCATGAACTG | qPCR |
| **GmMIR2111e_Rv** | GGAGGATATTGATTTTGTTGAACTGATG | qPCR |
| **GmMIR2111f_Fw** | GTAGCTGCTATCATACTAGACAGTTGA | qPCR |
| **GmMIR2111f_Rv** | CTTATATGAATTTCTCTACATGGATGGTC | qPCR |
| **Long SL miR2111_Rv** | GTCGTATCCAGTGCAGGGTCCGAGG TATTCGCACTGGATACGACTAAACC | Stem-loop reverse transcription |
| **Long SL miR1520_Rv** | GTCGTATCCAGTGCAGGGTCCGAGG TATTCGCACTGGATACGACTTGTCA | Stem-loop reverse transcription |
| **Mature miR2111_Fw** | CCGTTAATCTGCATCCTGA | Stem-loop qPCR |
| **Mature miR1520_Fw** | CGGACCATCAGAACATGACACG | Stem-loop qPCR |
| **miR2111 std curve_Fw** | CCGTTAATCTGCATCCTGAGGTTTAGTCGTA TCCAGTGCGAATACCTCGGACCCTGCACA | Standard curve construct |
| **miR2111 std curve_Rv** | GTGCAGGGTCCGAGGTATTCGCACTGGATA CGACTAAACCTCAGGATGCAGATTAACGGA | Standard curve construct |
| **SL universal_Rv** | GTGCAGGGTCCGAGGT | Stem-loop qPCR |
| **Artificial pre-miR2111_Fw** | ATAGTCTAATCTGCATCCTGAGGTTTACTTC TTTCTAAACCTCAGGATGCAGATTATC | miR2111 cloning |
| **Artificial pre-miR2111_Rv** | GAGCGATAATCTGCATCCTGAGGTTTAGAAA GAAGTAAACCTCAGGATGCAGATTAGA | miR2111 cloning |
| **L4440T7ter_Fw** | CCCCTGATTCTGTGGATAACCGTAT | Colony PCR/sequencing |
| **L4440T7ter_Rv** | CGGGCCTCTTCGCTATTACGC | Colony PCR |
| **Gm_miR2111a_Fw** | ATAGAATTCCTTTGAAGAATGATTGCTGGGTCC | Over-expression |
| **Gm_miR2111a_Rv** | GTAAAGCTTAGTGCTGGCAAGGAAGACG | Over-expression |
